# Bringing plants and soils to life through a simple role-playing activity

**DOI:** 10.1101/171009

**Authors:** Michael E. Van Nuland, Miranda Chen, Benjamin J. England

## Abstract

Interactions are at the core of many ecological and evolutionary forces in nature. Plant-soil interactions provide a rich example of the interconnectedness of living systems, but are hidden from everyday view and overshadowed in the classroom by more popular teaching examples involving animals, reptiles, or invertebrates. To highlight the importance and relevance of plant-soil relationships, we devised a simple role-playing activity suitable for college students. Specifically, the activity simulates how feedbacks between plants and soil environments influence plant species abundance and community richness. With this activity, students will gain a better understanding of these prolific, but overlooked, forms of biological interactions that impact the diversity and functioning of ecosystems.

## Motivation

Biological interactions are a predominant way in which students learn about the ecological and evolutionary processes that influence biodiversity. However, most general biology textbooks primarily use animals, reptiles, or invertebrates as case studies to demonstrate the importance of interactions in nature (Uno, 1994; Schussler *et al.*, 2010; Link-Perez *et al.*, 2010). This contrasts with the fact that (1) plants are ubiquitous, and students encounter them regularly in their daily lives, and (2) most interactions that plants rely on happen belowground. Since it can be difficult to present plants (and especially soils) in exciting ways, many students unintentionally cultivate a fauna-centric viewpoint of the natural world (Wandersee & Schussler, 2001). To highlight the importance and relevance of plant-soil relationships, we devised a simple role-playing activity suitable for college students.

## Context

Research on plant-soil interactions and their importance in ecology and evolution has blossomed in recent decades. Specifically, feedbacks occur when plants condition soil properties and, in return, are affected by the conditioned soils (Bever, 1994). Negative feedbacks reduce the performance of individuals of the same species relative to other species, resulting in negative frequency-dependent selection (Packer & Clay, 2000, Mangan *et al.*, 2010). Positive feedbacks encourage conspecifics to thrive in their respective soils more than heterospecifics, leading to the monodominance of single species (e.g., invasive species). These reciprocal interactions can shape the non-random assembly of plant communities (Bever 1994; van der Putten 2013).

## Purpose

The activity has two main purposes: (1) to engage students in a more active interpretation and discussion of the interactions between plants and soils, and (2) to connect these interactions to larger concepts of drivers of biodiversity and ecosystem function. This aligns with a core concept in Biology (*sensu* AAAS, 2011): living systems are interconnected and interacting. Plant-soil interactions provide a rich example of the interconnectedness of living systems, but are hidden from everyday view and overshadowed by more popular teaching examples. By actively role-playing plants and soils, students can see how these interactions operate in nature.

## Activity description

The primary learning objective of this activity is for students **to recognize how plant-soil interactions alter patterns of plant community diversity**. This activity simulates how plant-soil feedbacks influence species abundance and richness over time. It is recommended that the game first be played in **Negative Mode**. A **Positive Mode** variation is introduced at the end of this description.

**Negative Mode** demonstrates how negative plant-soil feedbacks promote and maintain diversity. When the same plant species and soil properties are matched, the plant dies. Oppositely, plants survive when species and conditioned soils are mismatched. For example, if a Spruce interacts with soil that has been previously conditioned by Spruce, that plant will die. But if the soil has been conditioned by Pine, Ash, or Ailanthus species, the Spruce will survive.

The directions for setup are as follows:

1. Split the class so that there are equal numbers of plant and soil players (Fig. 1a). Clear enough space in the room so that plant and soil players can stand in two opposing lines with no obstacles between the groups (Fig. 1b). Soil players each receive one blank notecard and paperclip. Include a pile of paper cutouts of the different plant species (3x the total amount of plant players for each species) to serve as the species pool, as well as a separate discard pile for when plants die and are not returned to the species pool.
2. Each plant player randomly draws one species from the species pool and returns to the line opposing soil players. Record the abundance of each plant species in the community at this initial time point (T_0_). These abundances are recorded on multiple graphs (one graph for each time point; Fig. 2a). Note: the game can be purposefully set at different levels of diversity at T_0_.

**Figure 1.**
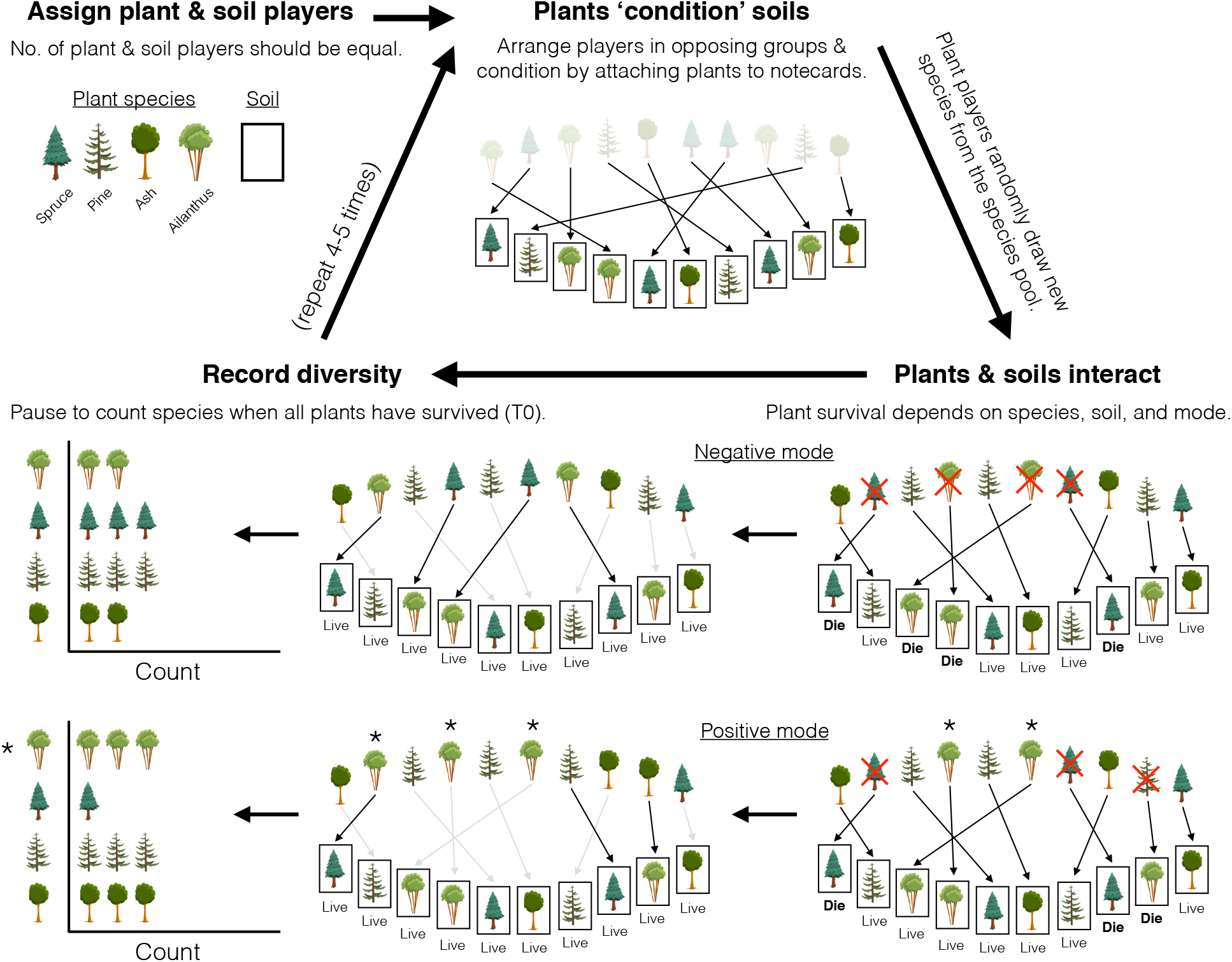
How to perform the activity. (**A**) The flow diagram outlines the general approach to the activity. The game is based on many random 1-on-1 interactions between plant and soil players. (**B**) The outcome of plant-soil interactions depend on the mode and identities of both plant and soil players. Plants that live remain matched with their soils (players stand next to one another; grey arrows). Plants that die discard their species, draw a new species from the species pool, and interact with any unmatched soil. In negative mode, plants die when they interact with soils conditioned by the same species (i.e., survival occurs when plants and soils are mismatched). In positive mode, select one plant species that will have positive interactions (e.g., Ailanthus species marked with asteriks). This species survives in all soil types, and no other species can survive in Ailanthus-conditioned soil. Plant species’ abundances are recorded once all plants are living to show how the species’abundance and community richness is shaped by plant-soil interactions.

After set-up, the gameplay begins:

(3) Plant players approach a random soil player to begin the conditioning phase that determines how plants change the physical, chemical, and biotic components of their soils based on traits related to their identity. For example, Ash leaves have lower carbon (C) to nitrogen (N) ratios than Spruce needles, which affects the quality of litter inputs to the soil and structures decomposer communities. To simulate conditioning, plant players hand their species to soil players, and soil players paperclip the species behind their soil card to hide the species that conditioned them.
(4) Plant players then randomly draw another species from the species pool, form a new line opposing the now-conditioned soil players, and interact with a random soil player. It is important that plant players do not know the conditioned status of soil players before interacting with them (just as tree seedlings cannot preferentially choose more hospitable soil locations in a forest). In this interaction, plants approach soils and show their species identity; in response, soils reveal to the plants what species they have been conditioned with. If they match, the plant dies.
(5) Plants that survive remain standing next to their respective soil player (no other plant can interact with soils that have a surviving plant). Plants that die discard their species in a separate pile (i.e., their genes do not get returned to the gene pool), before randomly re-drawing species from the species pool to interact with the remaining unmatched soil players. This gameplay continues until all plants are surviving and matched with a soil player. At this point, pause the game to record diversity with species’ abundances (Fig. 1b).
(6) Once diversity has been recorded, plants re-condition soils with their current species identity, making soil players clip the new species over their previous species before the next round (repeating Step 3). Again, the new conditioned statuses of soils should be hidden. Repeat steps 4-6 until you have measured diversity for multiple generations (T_0_–T_4_, or longer).

**Positive Mode** is a variation of the game with rules to show how positive plant-soil feedbacks are unstable and reduce diversity as one species becomes monodominant (e.g., invasion success through allelopathy). To play in Positive Mode, one plant species must be designated to have positive soil interactions, while all other plant species continue playing in Negative Mode. This designated species survives in all soil types conditioned by all species (including its own), and no other plant species can survive in soils conditioned by the designated species (Fig. 1b). For example, Ailanthus is an invasive tree species that produces toxic chemicals that inhibit the growth of nearby plants (Heisey 1990). Therefore, if Ailanthus were designated to have positive interactions, they would survive in any soil no matter what plant species conditioned that soil. In addition, Pine, Spruce and Ash species would die if they interact with Ailanthus-conditioned soil or their respective soils.

## Assessment, feedback, and suggestions

For a follow-up activity, we asked students to relate the direction of plant-soil feedbacks (positive or negative) to the abundance of specific plant species and total species richness (using results from the primary literature; Packer & Clay, 2000, Bais *et al*., 2003, Klironomos, 2002, Mangan *et al*., 2010, Bennett et al. 2017). Students successfully predicted that species with negative feedbacks would be rarer in communities that could sustain a greater number of total species, and that species with positive plant-soil interactions would be more abundant in less diverse communities.

We also asked the following question before and after the activity: “What do plant-soil feedbacks make you think of?”. Responses that included the words “fungi” or “mycorrhizae” increased 75%, and the word “diversity” appeared only in post-activity responses (7 out of 26 responses) (Fig. 2b). This suggests that students began to recognize how the nature of plant-soil relationships relate to biodiversity patterns.

**Figure 2.**
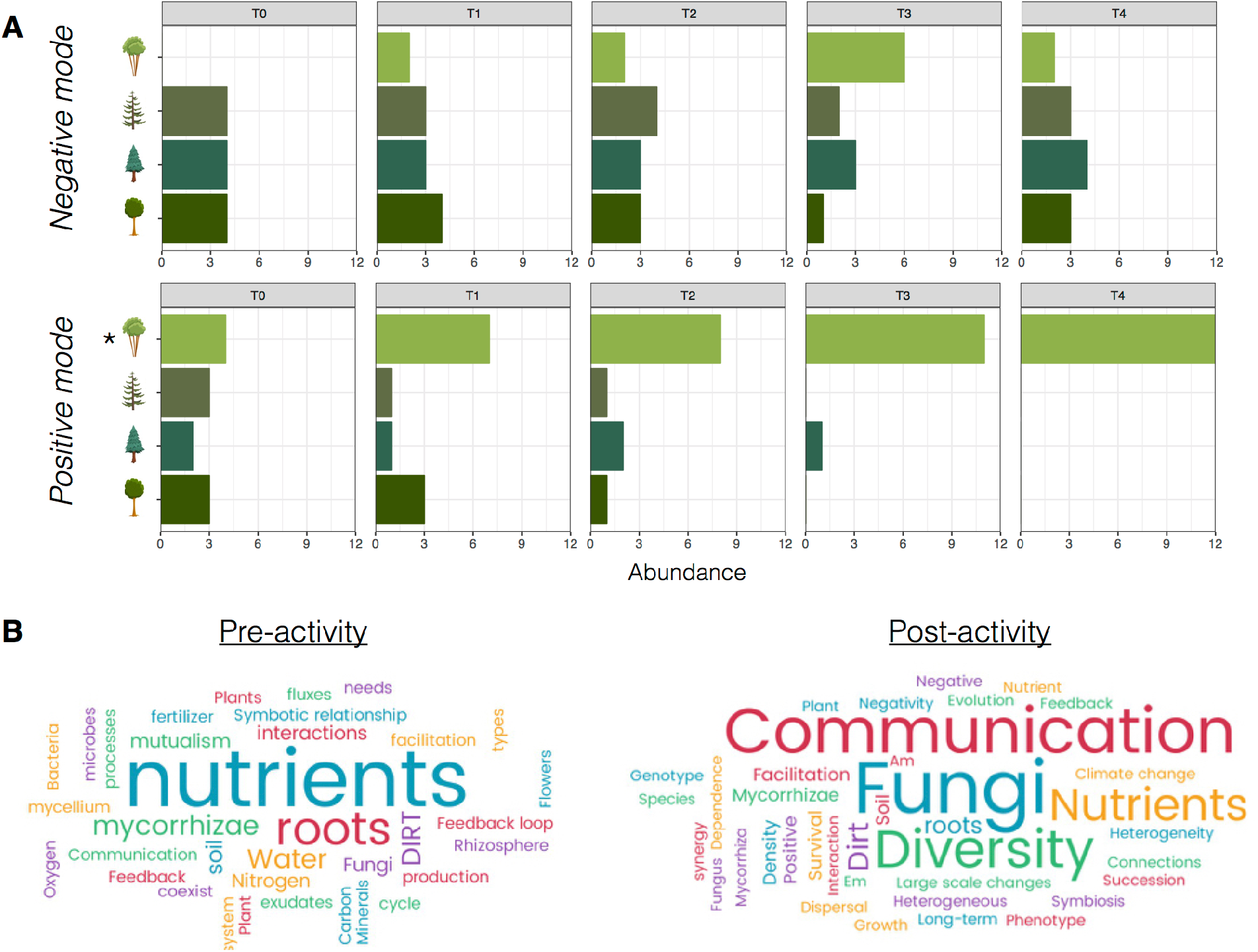
Evidence that the activity is successful and effective. (**A**) Modeling plant-soil interactions with the role-playing activity successfully depicted how negative interactions promote community diversity and stability, while positive interactions decrease diversity and disrupt communities. In negative mode, we purposefully set the plant community to have lower diversity (3 species) than the total number of species in the species pool (4 species). After one generation (T_0_-T_1_), students could see how negative plant-soil interactions increased plant diversity with the addition of Ailanthus into the community. In positive mode, plant community diversity abruptly declined as Ailanthus (an invasive tree species denoted with the asterisk) began conditioning a greater number of soil players that allowed them to persist and inhibited the survival of other species. Results are from a class of 24 undergraduates in an advanced ecology course (diversity was recorded on a whiteboard with the whole class at each time step and broad patterns were discussed when the activity ended). (**B**) Student responses when asked “What do plant-soil feedbacks make you think of?” pre- and post-activity and discussion. Word sizes reflect their total prevalence in the class responses. Overall, these responses show that the activity improved their understanding of how plant-soil interactions relate to patterns of biodiversity.

The activity takes ~30 minutes to complete and preceded a brief lecture and small group work in an upper-level Ecology course (24 students, 18-25 years old). Depending on student level and module topic, instructors using this activity could discuss a range of mechanisms, such as soil nutrient depletion by the plant, mutualistic benefits from mycorrhizal fungi, or build-up of soil-borne pathogens. Since plant-soil interactions have been explored in a variety of areas (van der Putten et al. 2013), the activity can be uniquely paired with different biology topics. For introductory students, shapes could be used in place of species to focus on the mechanics of feedback loops in nature.

This activity may be most applicable for small class sizes (20-40 students). In larger classes, this activity could be implemented as a demonstration with student volunteers or during discussion/laboratory sections. We found it best to use paper cutouts and notecards to drive home the role-playing aspects of the game. We have provided resources for teachers to print the species used in the current example (https://github.com/mvannuland/Species_supplies), but the activity is amenable to any suite of species (4-5 plant species is the appropriate number for a small class). With an inexpensive, time-efficient, and engaging activity, we hope to enable teachers to encourage student understanding of prolific, but overlooked, forms of biological interactions that impact the diversity and functioning of ecosystems.

